# Allochthonous groundwater microorganisms affect coastal seawater microbial abundance, activity and diversity

**DOI:** 10.1101/2023.05.14.540660

**Authors:** Keren Yanuka-Golub, Natalia Belkin, Nurit Weber, Meor Mayyani, Yehuda Levy, Itay J. Reznik, Maxim Rubin-Blum, Eyal Rahav, Yael Kiro

## Abstract

Submarine groundwater discharge (SGD) is a globally important process supplying nutrients and trace elements to the coastal environment, thus playing a pivotal role in sustaining marine primary productivity. Along with nutrients, groundwater also contains allochthonous microbes that are discharged from the terrestrial subsurface into the sea. Currently, little is known about the interactions between groundwater-borne and coastal seawater microbial populations, and their role upon introduction to coastal seawater populations. Here, we investigated seawater microbial abundance, activity and diversity in a site strongly influenced by SGD (*in-situ* observations), and through laboratory-controlled bottle incubations mimicking different mixing scenarios between SGD (either ambient or filtered through 0.1 µm/0.22 µm) and seawater. Our results demonstrate that the addition of <0.1 µm SGD stimulated heterotrophic activity and increased microbial abundance compared to control, whereas <0.22 µm filtration treatments induced primary productivity rates and *Synechococcus* growth. Amplicon sequencing of the 16S rRNA gene showed a strong shift from a SAR11-rich community in the reference SGD-unaffected coastal samples to a *Rhodobacteraceae*-dominated one in the <0.1 µm treatment, in agreement with their *in-situ* enrichment in the SGD field site. These results suggest that despite the significant nutrient input, microbes delivered by SGD may affect the abundance, activity and diversity of intrinsic microbes in coastal seawater. Our results highlight the cryptic interplay between groundwater and seawater microbes in coastal environments, which has important implications for carbon cycling and climate.

**Key Points:**

- Groundwater discharge into the coastal zone delivers both nutrients and allochthonous microbes.
- Groundwater microbes interact with seawater populations, by which affecting the delicate autotroph-heterotroph balance.
- Subterranean microbial processes are key drivers of food webs, potentially affecting biogenic carbon fluxes in the ocean and climate.

## 1 Introduction

Submarine groundwater discharge (SGD) is a globally important process, mixing the recirculated seawater inside the sediment and terrestrial groundwater flowing across the seabed with coastal waters (Burnett et al., 2003). This process of water exchange occurs at the subsurface land–ocean interface, termed subterranean estuary, STE (Moore, 1999; Rocha et al., 2021). Various dynamic driving forces, such as terrestrial hydraulic gradients, tides, waves, seasonal and interannual sea level may impact the chemical solute composition of SGD flux into the coastal ocean (Robinson et al., 2018). Thus, SGD acts as a major pathway for delivering terrestrial solutes across the land-ocean interface, and the STE is an important hotspot for biogeochemical reactions (Moore, 2010). This is especially true for ultra-oligotrophic coastal regions such as the southeastern Mediterranean Sea (SEMS), where the effects of the discharge are potentially most prominent (Rahav et al., 2020; Rodellas et al., 2015).

The terrestrial subsurface is one of the largest habitats for microorganisms on Earth (Griebler and Lueders, 2009), where biogeochemical processes, including the turnover of carbon and other nutrients, mineral cycling or pollutant degradation are primarily driven by diverse microbial populations (Probst et al., 2018). Generally, natural groundwater microorganisms live under completely dark conditions, with limited bioavailable organic carbon and phosphorous (Hofmann and Griebler, 2018); thus, they are forced to develop diverse metabolic and physical strategies to survive (Griebler and Lueders, 2009). Recently, *in situ* dark inorganic carbon fixation rates were reported in a carbonate aquifer (Overholt et al., 2022). Estimations were similar to those measured in oligotrophic marine surface waters, indicating chemolithoautotrophs play an important role in subsurface ecosystems processes. Contrary to the relatively stable conditions maintained in inland aquifers, coastal aquifers entail rapid fluctuations and steep physiochemical gradients to which STE microbes need to quickly adapt. This results in a highly diverse microbial community composition along the hydrological continuum of SGD sites, reflecting a mixture of seawater and groundwater-associated taxa (Adyasari et al., 2019; Chen et al., 2020; Degenhardt et al., 2020), as well as microbial functional groups that follow redox gradients (McAllister et al., 2015; Purkamo et al., 2022). Therefore, ubiquity of diverse prokaryotic groups in coastal groundwater ecosystems that are metabolically-active (Miyoshi et al., 2005), or dormant due to scarcity of nutrients and carbon sources in the subsurface habitat (Hofmann and Griebler, 2018) could potentially affect bacterioplankton communities upon discharge into the marine environment (Lee et al., 2017; Ruiz-González et al., 2021).

Bacterioplankton community variation across estuarine gradients were shown to be driven primarily by salinity (Fortunato and Crump, 2011; Herlemann et al., 2011). Adyasari et al. (2020) also found that salinity was the most decisive variable that shaped the microbial community composition across surface water samples, while dissolved nitrogen and phosphorus were the major predictor of community shift within subsurface water samples. At the same study site, higher microbial diversity was found in coastal porewater in comparison to surface water samples, consisting of a diverse mixture of fresh water and marine microbes. The presence of microorganisms transported from external sources that are hydraulically connected show a bidirectional dispersal of groundwater and marine species (Ruiz-González et al., 2021). It still remains unknown, however, whether the transported communities tolerate such wide ranges of conditions to remain active. Nevertheless, at a different study (Adyasari et al., 2019), brackish porewater samples were more similar to freshwater samples than to saline samples, comprising mainly taxonomic groups associated with nitrogen transformation (nitrification and denitrification) in natural water systems. Given that nitrogen cycling is a key pathway in the subterranean estuary (Erler et al., 2014; Hays and Ullman, 2007; Slomp and Van Cappellen, 2004), these findings indicate the possibility of active biological transformation of dissolved nitrogen along the land-water interface.

SGD often exceeds the Redfield ratio and alleviate co-limitation of N and Si in marine environments (Santos et al., 2021). The examined effect is mostly attributed to the delivery of dissolved nutrients stimulating the growth of bloom-forming phytoplankton (Garcés et al., 2011; Lecher et al., 2017, 2015). However, supplements of groundwater to seawater is not always straightforward and behave in a dose-depended response. For example, Lecher et al., (2015) observed through incubation experiments that 10% groundwater addition rather than 20-50% (volumetric) yields the highest positive effect on chlorophyll a concentrations and phytoplankton abundance, despite the fewer nutrients introduced. Contrary, lower addition volume ratios (1-10%) did respond at a dose-dependent manner (Lecher et al., 2015). The authors concluded that these responses were attributed to the elevated nutrients supplied by the SGD, but this was achieved only to a certain level. This implies a direct biological effect of SGD through the introduction of subsurface bacterial cells into the coastal environment, which could potentially affect the native coastal microbiome. Recently it has been reported that STEs are continuously subjected to a bidirectional inoculation of microbial taxa from the marine and fresh groundwater endmembers (Adyasari et al., 2019). However, investigations mainly focused on health implications of the marine environment due to the transport of pathogenic or fecal indicator bacteria through coastal aquifers or sediments, neglecting the ecological implications. Currently, the role of coastal aquifers as sources of microbial diversity and the responses of marine microbial communities to groundwater-derived microorganisms is still unknown.

The occurrence of specific heterotrophic bacterial groups associated with cyanobacteria is dynamic and highly influenced by biotic and chemical factors. For example, Garcés et al., (2011) found in the Mediterranean Sea that the addition of groundwater to marine communities resulted in *Synechococcus* fast growth, due to the nutrient addition. But, this observation was not consistent in other coastal settings, where other phytoplankton groups dominated (Chamberlain et al., 2014). Marine *Synechococcus* are ubiquitous cyanobacteria in the global ocean and is abundant in both estuarine and coastal waters (Dufresne et al., 2008; Huang et al., 2012). Synergistic interactions between *Synechococcus* and alphaproteobacteria heterotrophic bacteria (*Rhodobacteraceae*), gammaproteobacteria (*Alteromonadaceae*) (Dang and Lovell, 2016; Wang et al., 2021; Zheng et al., 2018), and the Cytophaga-Flavobacteria-Bacteroides (CFB) group are fundamental in the marine food web (Buchan et al., 2014). In oligotrophic surface oceans, the three bacterial groups *Prochlorococcus*, marine *Synechococcus*, and heterotrophs in the SAR11 clade often represent more than 50% of free-living bacterial cells (Becker et al., 2019). The abundant SAR11 is a strict oligotrophic heterotroph (Kearney et al., 2021), thus other heterotrophic groups that are adapted to higher organic carbon concentrations such as those associated with phytoplankton blooms are often favored (Buchan et al., 2014; Tada et al., 2011; Teeling et al., 2012). Previous studies showed that the *Roseobacter/Rhodobacter* group of bacteria may respond as generalists because they remain active regardless of large fluctuations in organic matter, whereas the *Alteromonas* group are defined as specialists for using organic matter and became abundant only in the presence of phytoplankton blooms (Tada et al., 2011). Given the delicate autotrophic-heterotrophic metabolic balance in coastal marine ecosystems, SGD is an important factor that should be accounted for with respect to its chemical and biological effects on the ocean.

We recently compared a site strongly influenced by SGD (Achziv, northern Israel) and a nearby nutrients-poor reference site at the oligotrophic Israeli shallow rocky coast (Rahav et al., 2020). At the SGD site (Achziv), we showed that SGD contributes to high concentrations of dissolved nitrate and silica in comparison to the reference coastal seawater, resulting in elevated *in-situ* phytoplankton biomass and primary productivity. The main objective of this study is to further elucidate the response of marine heterotrophic bacterial taxonomic and functional responses to SGD by addressing the following questions: i) Do coastal microbial communities respond to the nutrient-rich source of SGD in a dose-dependent manner (i.e. higher SGD-enriched mixtures correspond linearly with higher microbial activity and abundance)? ii) How do discharged groundwater microbes shape the activity and composition of the coastal microbiota? To explore these questions, three complementary incubation experiments were designed to illustrate the taxonomic and physiological response upon introducing groundwater and coastal communities. Additionally, porewater samples during three field campaigns were collected to follow composition, abundance and activity of the natural microbial communities at the Achziv SGD site. Our results show that groundwater prokaryotes transported through SGD influence coastal microbial activity and diversity, resulting in different heterotrophic composition patterns. We were able to mimic a similar pattern in bottle incubation experiments by filtering the groundwater prior mixing with coastal seawater. This points at potential competitive interactions for available nutrients in oligotrophic marine environments.

## 2 Materials and Methods

### 2.1 Field site, sampling procedures, and sample analyses

**Field site**: Groundwater and seawater samples were collected from the shore sediments and surface seawater at Achziv (Lat. 33°3’52 N, Lon. 35°6’14.94 E), located along the Northern Israeli Mediterranean shoreline (Figure 1). Achziv coastal aquifer structure is comprised of Pleistocene calcareous sandstone, beach rocks, and carbonate sand, providing a complex groundwater flow system. About five km east of the coastline, the carbonate Galilee Mountains (Judea Group) are elevated. The Quaternary permeable filling of the coastal aquifer is recharged mainly from the east, by precipitation on these carbonate mountains (Kafri and Kessler, 2001; Paldor et al., 2020). Previous studies show fresh SGD (or a fresh SGD component) both shallow along the shoreline (Weinstein et al., 2006) and in deeper parts, through preferential flow in the Judea Group aquifer (Paldor et al., 2020, 2019). The exact SGD sampling locations were selected based on the density and electric conductivity to identify the fresh groundwater and fresh-saline transition zone and to track changes in the salinity distribution compared to previous seasons.

**Figure 1:**
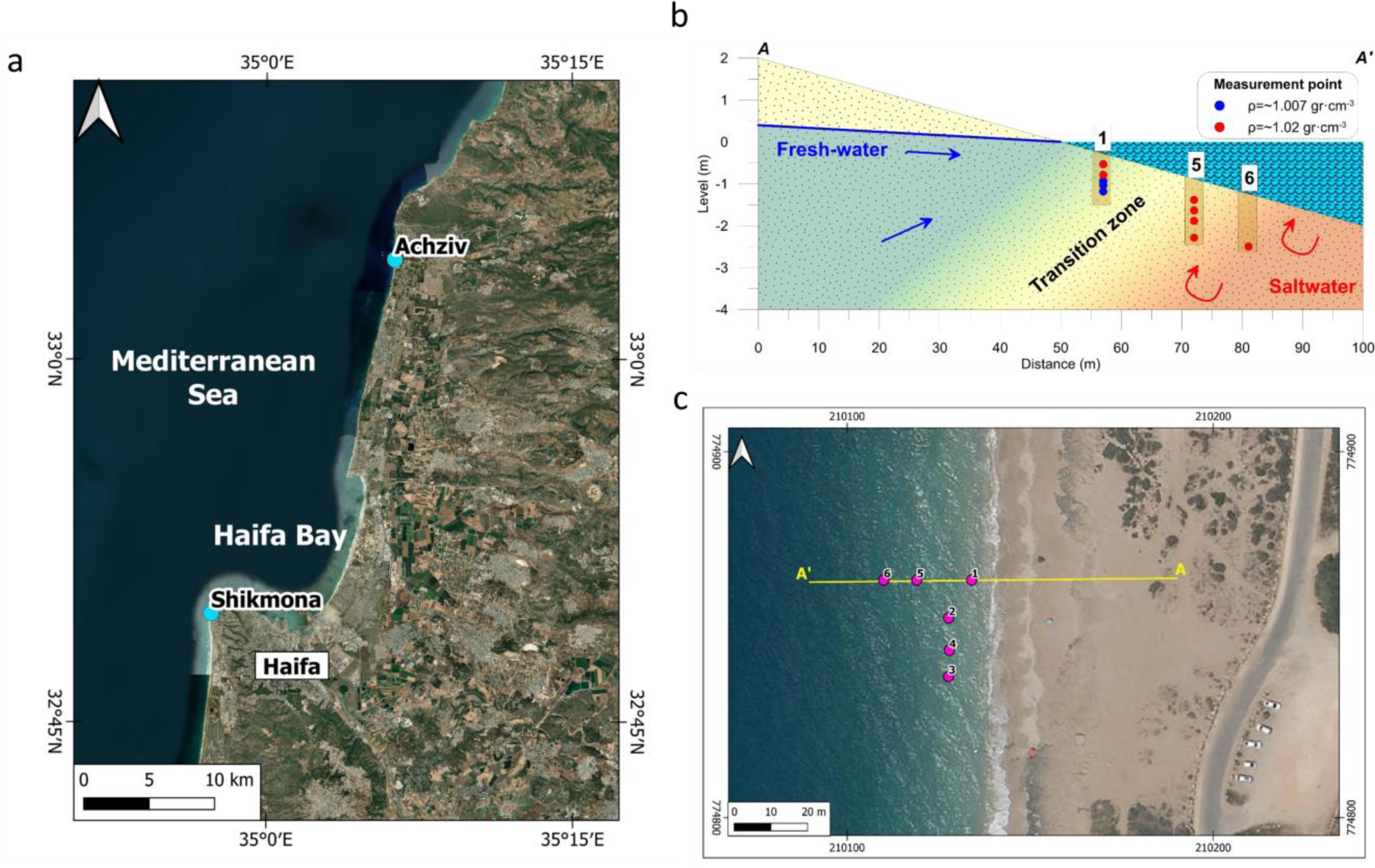
(a) Map showing the location of the two sampling areas at the Israeli coastline in the SE Mediterranean Sea: Achziv, the SGD-site (33° 3′52 N, 35° 6′14.94 E) and Shikmona (32° 49′34 N, 34° 57′20 E) within the Haifa Bay (sampled as the reference non-SGD site during the incubation experiments). **(b)** Schematic representation of the field site in Achziv showing the location of the piezometers (brown bars, numbers indicate piezometer ID). Piezometer samples are referred to in the text as either “Low or High salinity porewater” samples (n = 9 and 13, respectively) and are distinguished from the surface seawater samples (n = 11). Accordingly, red dots indicate high salinity (∼30 ppt) porewater and blue dots indicate low salinity porewater samples (∼17 ppt). **(c)** aerial photography with the piezometers setup for Achziv sampling campaigns. The numbers indicate piezometer ID along a perpendicular (1, 5-6) and parallel (2-4) to the shoreline cross sections.

**Sample collection**: The fieldwork included three campaigns at Achziv Beach, Israel (August 2020, February and July 2021). Each field campaign lasted 2-5 days and covered at least 2 tidal cycles. Physical parameters were measured, and porewater samples were collected on the shoreline using piezometers (AMS piezometers that reach depths of <2 meters) and a portable peristaltic pump (Masterflex®, Cole-Parmer, Germany). The density (g cm^−3^), electric conductivity (mS/cm), temperature (°C) and pH, of surface seawater, porewater and groundwater were measured on-site at the time of the sampling. Based on average temperature and water density measurements, salinity (ppt) was calculated.

Samples for microbial analysis were collected from the piezometers and divided to aliquots: First, for community analysis, samples were immediately filtered through polycarbonate 0.2 μm pore size filters (Merck, Israel), which were kept on ice and transported to the laboratory on the same day. Filter samples were stored frozen (−20°C) until DNA extraction. Second, for Pico-/nano-phytoplankton and heterotrophic prokaryotic abundance, non-filtered samples were chilled on ice and transported to the laboratory on the same day.

### 2.2 Bottle incubation experiments

Three incubation experiments were conducted to determine the contribution of SGD to the coastal microbial community. The experiments were conducted with five different treatments (including no-mixing control of reference coastal seawater, i.e., ambient seawater that is not exposed to SGD) in triplicates as described in Table 1. The first experiment (Exp. 1) was designed to test the relative contribution of brackish discharged groundwater (salinity = 7.93 ppt) on the microbial productivity and abundance of reference coastal seawater by mixing different ratios (1, 5, 10 and 20%) of discharged groundwater to reference coastal seawater. Discharged groundwater was collected into acid-cleaned containers on the day the experiment initiated near Achziv National Park (33° 3′52 N, 35° 6′14.94 E). At this sampling site, a significant groundwater discharge was spotted (Rahav et al., 2020; Weinstein et al., 2006). The second and third experiments (Exp. 2; Exp.3) were designed to extend Exp. 1 and aimed to specifically investigate how groundwater-derived microorganisms affect the activity and abundance of marine organisms once discharged into the sea. For these experiments, fresh groundwater (FGW) was collected from drilling wells and pumped into 20 L acid-cleaned sample-rinsed carboys the same day the experiment was initiated. At the laboratory, fresh groundwater was either filtered through a 0.1 μm polycarbonate filter (Exp. 2) or serially filtered through 0.22 and 0.1 μm polycarbonate filters (Exp. 3). The filtrate was added to seawater in different mixing scenarios as described in Table 1.

**Table 1:** Summary of the three-bottle incubation experimental designs

|  | Experiment aim | Groundwater source | Control groups<br>No addition |  | Treatment groups<br>Mixing ratios with ambient seawater<br>(v:v)* |  |  |  |
| --- | --- | --- | --- | --- | --- | --- | --- | --- |
| **Exp. 1** | Investigate dose-dependent responses | Brackish discharged groundwater (Achziv site) | Reference ambient seawater |  | 1% | 5% | 10% | 20% |
| **Exp. 2** | Compare filtered fraction supply of dissolved inorganic nutrients, to ambient groundwater | Inland fresh groundwater | Reference ambient seawater | Fresh groundwater | 5% | 5% of 0.1 $\mu\text{m}$ filtered FGW | | |
| **Exp. 3** | Investigate sequentially filtered groundwater and dilution effect (DDW) | Inland fresh groundwater | Reference ambient seawater | | 5% | 5% DDW | 5% 0.1 & 0.22 $\mu\text{m}$ filtered FGW | |

\* For all incubation experiments, the % ratio indicates adding groundwater to ambient seawater, unless stated otherwise.

Ambient coastal seawater was collected by pumping at the Israel Oceanographic and Limnological Research Institute (IOLR) into acid-cleaned carboys, and mixed with either brackish groundwater (Exp.1) or fresh groundwater (Exp. 2, Exp. 3) at the desired ratios and filtration size (Table 1). Then, each mixture was randomly distributed into acid-washed transparent polycarbonate Nalgene bottles (4.5 L for Exp. 1 and 2; 250 ml bottles for Exp 3).

The duration of the experiments was 3-5 days, and samples were taken for the following analyses: chlorophyll *a* (Exp. 1 & 2, every 24Hr.), dissolved nutrient concentrations (Exp. 2 & 3 T zero and T final), flow cytometry (bacterial and phytoplankton abundance, every 24Hr.), primary and heterotrophic production rates (Exp. 1 & 2, every 24Hr.; Exp. 3 T zero and T final), methods are specified below. All results were normalized to the dilution factor (for example, in the 20% treatment group, all measurement values were multiplied by 1.2). All bottles were incubated in a flow-through tank at the IOLR through which ambient Eastern Mediterranean water was continuously pumped to maintain surface ocean temperature. The incubation tanks were covered with an illumination net to maintain natural light (representing a full-day cycle).

### 2.3 Heterotrophic prokaryotic and Pico-/nano-phytoplankton abundance

Samples (1.8 mL) were fixed with glutaraldehyde (final concentration 0.02 % v:v, Sigma-Aldrich G7651), frozen in liquid nitrogen, and later stored at −80°C until analysis. The abundance of autotrophic pico- and nano-eukaryotes, *Synechococcus* and *Prochlorococcus*, and other heterotrophic prokaryotes (bacteria and archaea) was determined using an Attune® Acoustic Focusing Flow Cytometer (Applied Biosystems) equipped with a syringe based fluidic system and 488 and 405 nm lasers. To measure heterotrophic prokaryote abundance (hereafter refer to as heterotrophic abundance), a sample aliquot was stained with SYBR Green (Applied Biosystems).

### 2.4 Heterotrophic productivity

Prokaryotic (bacteria and archaea) heterotrophic production was estimated using the ^3^H-leucine incorporation method (Perkin Elmer, specific activity 100 Ci mmol^−1^). Water samples (1.7 mL each) were incubated in the dark (wrapped in aluminum foil) with ∼ 10 nmol hot leucine L^−1^ for 4 h (Rahav et al., 2019). For control treatments, an additional sample was immediately added with 100 µL of 100 % trichloroacetic acid (TCA, 4^∘^C) along with ^3^H-leucine, and was carried out in triplicates. The incubations were terminated with TCA and were later processed following the micro-centrifugation technique (Smith and Azam, 1992) and added with 1 mL of scintillation cocktail (Ultima-Gold). The samples were counted using a TRI-CARB 2100 TR (Packard) liquid scintillation counter.

### 2.5 Primary productivity

Photosynthetic carbon fixation rates were estimated using the ^14^C incorporation method (Nielsen, 1952). Briefly, collected water samples (50 mL) were immediately spiked with 5 µCi of NaH^14^CO3 (Perkin Elmer, specific activity 56 mCi mmol^−1^). The samples were incubated for 24 h under in situ natural illumination. The incubations were terminated by filtering the spiked seawater through GF/F filters (Whatman, 0.7 µm pore size) at low pressure (∼ 50 mmHg). Measurements for dark controls were also performed and were removed from the sample reads. The filters were placed overnight in 5 mL scintillation vials containing 50 µL of 32 % hydrochloric acid to remove excess ^14^C, after which 5 mL of scintillation cocktail (Ultima-Gold) was added. Radioactivity was measured using a TRI-CARB 2100 TR (Packard) liquid scintillation counter.

### 2.6 Inorganic nutrients determination

Samples from the field were immediately filtered through 0.2 μm pore size filters into acid-washed plastic vials, chilled on ice and transported to the laboratory on the same day. Samples were stored frozen until analysis. Experimental samples at time-zero and final time point were filtered through 0.2 μm pore size filters into acid-washed plastic scintillation vials, immediately frozen (− 20°C) and kept frozen until analysis. Dissolved nutrients (NO_2_ + NO_3_ = NOx, PO_4_, and Si(OH)_4_) were analyzed in the Interuniversity Institute for Marine Sciences (IUI, Eilat) using QuikChem 8000 flow injection analyzer (Lachat Instruments, Milwaukee, USA). The measurement is based on a color reaction created by each of the nutrients with its unique reagent to create a color complex with a wavelength in the visible light range, which is absorbed by the device’s spectrophotometer.

### 2.7 DNA extraction, amplicon sequencing, and analysis of bacterial community

Seawater or groundwater (4-5 L) were filtered using a peristaltic pump onto polycarbonate membrane filters (0.1 or 0.2 µm, 47 mm, Merck, Israel) and stored at −20°C until extraction (n=1 per sample except in selected samples where n=4). After thawing, each filter was cut into small pieces using a sterile scalpel blade, which was placed immediately into PowerSoil DNA bead tubes and extracted with the dNeasy PowerSoil Kit (Qiagen, USA) following the standard protocol. To disrupt the cells, the FastPrep-24™ Classic (MP Biomedicals, USA) bead-beating was used (2 cycles at 5.5 m sec–1, with a 5 min interval).

To generate 16S rRNA gene libraries, the V3–V4 hypervariable region of the 16S gene was amplified (∼ 465 bp) using the 341F/806R primers containing CS1 and CS2 linkers – ACACTGACGACATGGTTCTACACCTACGGGNGGCWGCAG and TACGGTAGCAGAGACTTGGTCTGG ACTACHVGGGTWTCTAAT. PCR conditions were as follows: initial denaturation at 95°C for 5 min, 28 cycles of denaturation (95°C for 30 sec), annealing (50°C for 30 sec) and extension (72°C for 60 s). Sequences were obtained on the Illumina MiSeq platform in a 2 × 250 bp paired-end run (HyLabs Israel). Demultiplexing and assignment of the amplicon sequence variants (ASVs) were performed using the Qiime2 (v. 2020.2) (Bolyen et al., 2019) platform, and DADA 2 was used as a quality control for chimeric reads (Callahan et al., 2016). The two 250-bp paired-end sequences were merged to obtain a single read (approximately 416.27 bp, mean length). Taxonomic assignment of the ASVs was done with q2_feature_classifier (Bokulich et al., 2018) using the Silva database (release 138-99) for 16S rRNA gene sequencing at 99% identity (Quast et al., 2012).

Alpha diversity and variance calculations were performed using Mothur (v.1.48.0). ASV data table was filtered (removing ASVs that appear in total <10 times) and repeatedly subsampled to the lowest number of reads among all samples (12,538 reads x 100 times). Diversity indices included Invsimpson, Shannon and Chao. Beta diversity was analyzed using Bray–Curtis metrics of community dissimilarity and plotted using pCoA ordination with PAST v4.09 (Hammer et al., 2001). Analysis of similarity (ANOSIM) (9999 permutations) was used to make pairwise comparisons between the different samples’ types (Field: Low and high-salinity porewater, Achziv surface seawater and experimental treatment groups: control ambient coastal seawater (SGD-non influenced), non-filtered, and 5% filtered 0.1μm pore size) treatments.

## 3. Data Analysis

### 3.1 Statistical analyses

Statistical analysis was performed in GraphPad Prism 5.03 (GraphPad, La Jolla, CA, USA). One-way analysis of variance (ANOVA) was used to determine statistical differences between the control and different mixing treatments for the three-bottle incubation experiments, and the three field campaign sample types. Multiple pairwise comparisons were achieved using Tukey 95% confidence intervals.

### 3.2 Accession numbers

Raw data from Illumina MiSeq sequencing are deposited to the National Center for Biotechnology Information (NCBI) Sequence Read Archive (SRA) under BioProject number PRJNA973031.

## 4 Results and Discussion

### 4.1 Groundwater discharge enhances coastal microbial abundance but constrains activity

Three bottle incubation experiments were designed to independently study the microbial response of groundwater additions (brackish or fresh) to ambient coastal seawater. Specifically, the aim of the first experiment (hereafter, Exp. 1) was to investigate the response of ambient coastal microorganisms (unexposed to SGD and corresponding microbes) to SGD by additions of discharged groundwater at pre-determined volumetric percentages (1%, 5%, 10%, 20%) relative to ambient seawater (Figure 2). The chosen volumetric range aimed to demonstrate naturally occurring conditions that could occur in an SGD site concerning point-source discharge flux. Temporal measurements indicated that a ∼40-Hr incubation period yielded maximal production rates, while longer times resulted in saturation of heterotrophic and primary production rates (Supplementary data, S1). Thus, the following results are presented for the 42-Hr incubation time point. Results showed that heterotrophic prokaryotic abundance increased in a dose-dependent manner, ranging from 4.8×10^5^±0.1×10^5^ cells ml^−1^ in the control to 19×10^5^±1.4×10^5^ cells ml^−1^ in the 20% GW treatment (Figure 2A). Contrary, heterotrophic production rates reached a plateau already at 1% GW enrichment; 11.8±2.4 µg C L^−1^ day^−1^, while the control values were ∼6 fold lower (Figure 2B). Primary production rates (Figure 2C) and chlorophyll *a* concentrations (Supplementary data, S1), reached a plateau at 5% treatment, in agreement with an earlier study from the intertidal area affected by SGD in Monterey Bay (Lecher et al., 2015).

**Figure 2:**
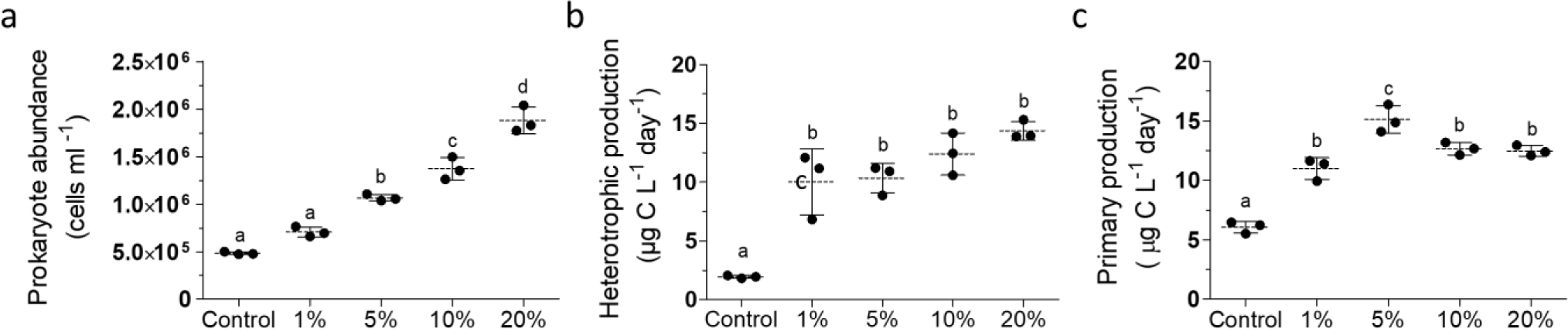
Scatter dot plots of surface seawater (control) prokaryote abundance (a), heterotrophic production rate (b) and primary production rate (c) following dilution with discharged brackish groundwater (1-20% v:v) or un-amended seawater (control). Results are shown for the 2^nd^ time point of the experiment (42hr) as described in the text. Lowercase letters indicate significant differences between treatments (using ANOVA followed by Tukey posthoc tests, p ≤ 0.05). The temporal variability of all time points is shown in Supplementary Information (S1). The dilution factor was calculated for each treatment to account for the volume of groundwater added to ambient seawater.

These observations suggest a further interaction between the groundwater microbes and the seawater microbes, which limits activity, but not abundance, despite the addition of nutrients.

### 4.2 Groundwater microbes transported through SGD may limit coastal production rates

To assess the microbial contribution of SGD to heterotrophic production, a second experiment was designed (hereafter, Exp. 2, Table 1), in which filtered (<0.1 μm) and non-filtered (5 and 20%) groundwater samples were mixed with the reference ambient coastal seawater. Amendment of filtered groundwater presented primarily the addition of dissolved nutrients, whereas non-filtered groundwater supplied nutrients along with endogenous groundwater microbes. Temporal measurements indicated that a ∼67-Hr incubation period yielded maximal production rates (Supplementary data, S2). Thus, the following results are presented for the 67-Hr incubation time point (Figure 3, Supplementary Table S1). Natural SGD-affected coastal measurements from Achziv were also evaluated (samples were collected at two different seasons: February 2021 and June 2022, Table 2). Compared to non-filtered groundwater and the control treatments, the filtered treatment showed the highest change for prokaryote abundance (Figure 3A) and heterotrophic production rates (Figure 3B). While prokaryote abundance of Achziv surface seawater was similar to the filtered treatment, heterotrophic production was significantly lower (similar to non-filtered treatments and control). Contrary, no changes in primary production rates were observed between the different groundwater amendment treatments (Figure 3C). These rates were significantly higher than the control (ANOVA and Tukey posthoc tests, p ≤ 0.05), but slightly lower than Achziv surface seawater. These findings lie in agreement with the results of experiment 1, where prokaryote abundance is dose dependent and activity reaches a stable threshold, supported by the natural Achziv measurements (Figure 3). Furthermore, changes in concentration of phosphate (PO_4_), silica (Si(OH)_4_), nitrate (NO_3_), and ammonium (NH_4_) over the incubation experiment period indicated that mainly PO_4_ and Si(OH)_4_ were uptaken by the microbial community provided by groundwater amendments (Supplementary data, S3). The 20% non-filtered treatment utilized significantly more of these nutrients than the other filtered and non-filtered treatments (Supplementary data, S3), as observed in a previous study (Lecher et al., 2015).

**Figure 3:**
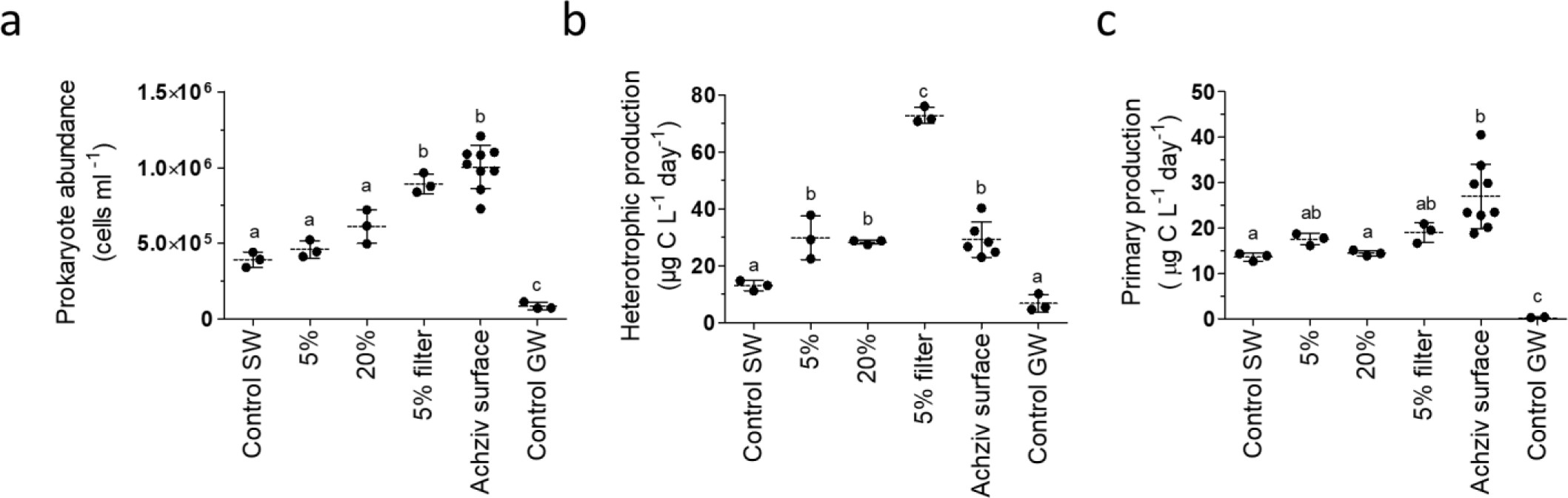
Scatter dot plots of surface seawater prokaryote abundance (a), heterotrophic production (b) and primary production (c) following dilution with fresh groundwater (5% and 5% filtered 0.1 μm v:v), un-amended seawater (control SW) or fresh groundwater (Control GW) during Experiment 2. Results are shown for the 3^rd^ time point of the experiment (67hr) as described in the text. Surface seawater measurements from Achziv (SGD-site) were also added to this analysis. Lowercase letters indicate significant differences between treatments (using ANOVA followed by Tukey posthoc tests, p ≤ 0.05). The temporal variability of all time points is shown in the Supplementary Information (S2). The dilution factor was calculated for each treatment to account for the volume of groundwater added to ambient seawater.

**Table 2:**
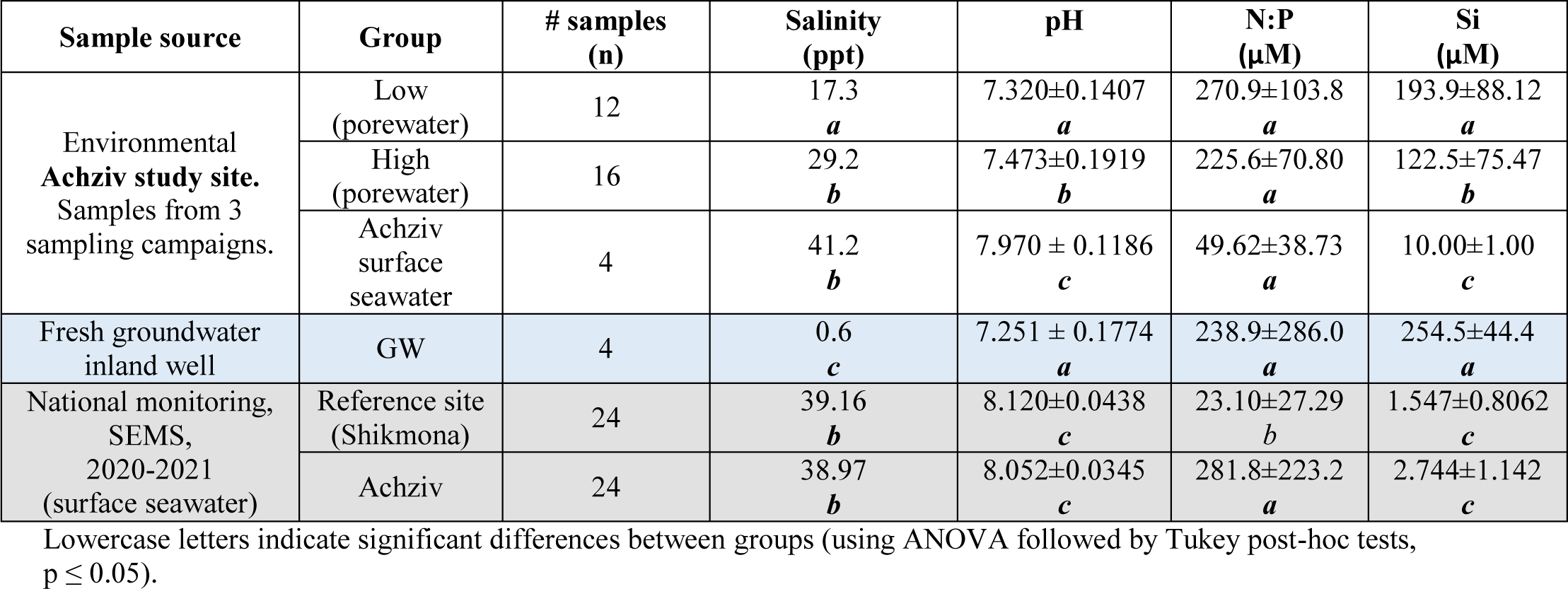
Summary of environmental parameters averaged by sample groups

The higher increase in prokaryote abundance or heterotrophic production in the filtered vs. the non-filtered addition suggests that groundwater microbes affect the seawater communities and potentially outcompete them for the added nutrients, highlighting the intricate interactions between autotrophic and heterotrophic microbes in oligotrophic realms such as the Mediterranean coast.

The control fresh groundwater (collected from an inland aquifer), incubated without mixing, showed the lowest values of abundance and productivity (Figure 3). While lower phototrophic abundance and activity are negligible, as expected in subsurface environments, the lower prokaryotic abundance (8.6±2.4×10^4^ cells/ml) and heterotrophic productivity (6.74±3.0 µg C L^−1^ day^−1^) are in agreement with values reported for pristine inland fresh groundwater. These observations show that a large fraction of communities from inland aquifers are locally inactive (Griebler and Lueders, 2009). Despite their low numbers and activity rates, incubation experiments of groundwater communities showed that groundwater bacteria can respond rapidly to environmental changes as implied from our data as well, and grow even when mixed with marine waters (Adyasari et al., 2019). This could ultimately shape coastal microbial diversity and activity.

### 4.4 Groundwater microbes transported through SGD may shape coastal community composition

Three sampling campaigns (August 2020, February 2021 and July 2021) were conducted at a field site, highly influenced by SGD (Achziv, northern Israel). Porewater samples were collected on the shoreline using piezometers at different depths (0.25-1m, Figure 1). According to the recorded porewater density values, the porewater samples were divided into two groups: low and high salinity. In addition, we sampled surface seawater (Table 2). The measured parameters of these samples were compared to values obtained for both Achziv and the reference site (Shikmona) during the routine monitoring program by Israel Oceanographic and Limnological Research, IOLR (Table 2).

The measured parameters reported for Achziv surface seawater during the three field campaigns were comparable to the ongoing monitoring results obtained during the years 2020-2021 (salinity, pH, and Si(OH)_4_, ANOVA, p > 0.05) with no significant seasonal or annual differences. As expected, the low-salinity porewater samples showed significantly lower salinity/pH values compared to high-salinity porewater and surface water (Table 2), further supporting the contribution of freshwater inputs. The mean N:P ratio of both high- and low-salinity porewater samples were comparable to values obtained during routine monitoring of the Achziv site for the last two years. The N:P values obtained at all Achziv samples were significantly higher than in the Shikmona reference site, signifying the influence of SGD as an important nutrient source. Notably, Si(OH)_4_, PO_4_ and NO_3_ concentrations of porewater samples exceeded the surface seawater samples by an order of magnitude (Table 2), and lie on a conservative mixing line with water salinity (Supplementary data, S4). Higher PO_4_ concentrations were observed in all Achziv porewater samples compared to the reference site, ranging from 1.0–3.7 μM, but generally peaked at the lower-salinity samples (2.9±0.6 μM).

Sequencing of the 16S rRNA gene resulted in 1,333,846 quality sequences, which clustered into 2235 ASVs, including field campaigns and bottle incubation experiments. The number of ASVs per sample varied between 36 and 493. Microbial community diversity of samples collected in the Achziv field site (low and high-salinity porewater and surface seawater) was compared with those obtained from the experimental treatments, namely control seawater (ambient seawater, not influenced by SGD), non-filtered treatments (1%, 5%, 20%), filtered (5% 0.1 µm pore size) and fresh groundwater from inland wells. Among these, the low-salinity porewater group was the most diverse for the three alpha diversity indices investigated (Figure 4), indicating a high heterogeneity of the low-salinity microbial community, and a relatively even community inferred from the higher inverse Simpson index (Figure 4c).

**Figure 4:**
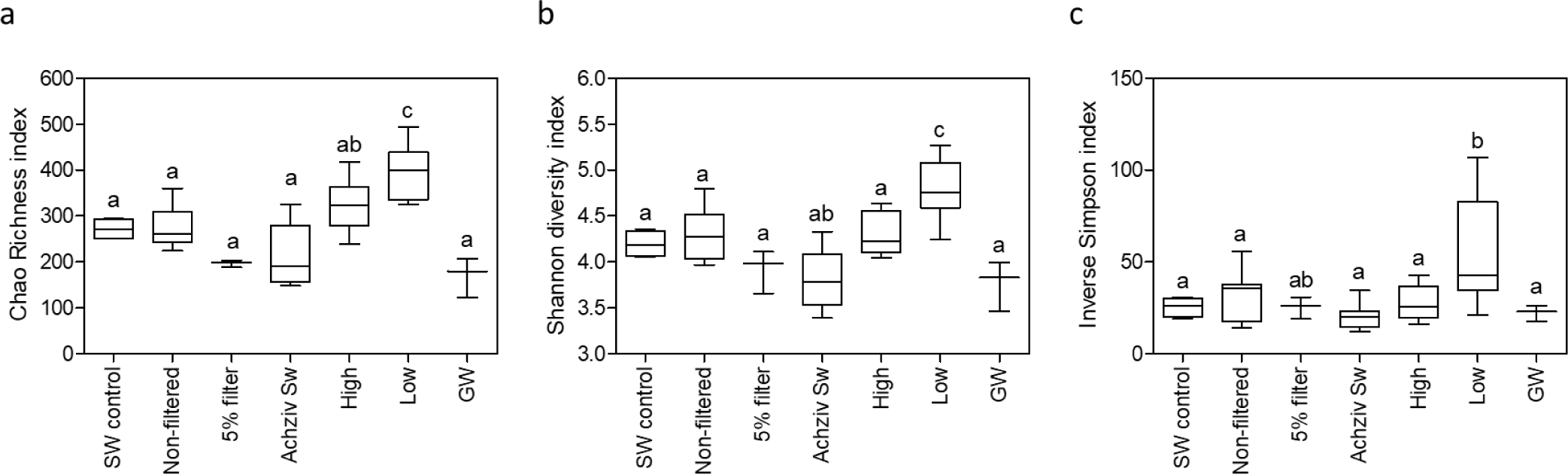
Box-Whisker plots showing the interquartile range (25th to 75th percentile) and the median value (horizontal line within the box) of the microbial communities’ alpha diversity grouped according to field and experimental samples assessed by the Chao richness estimating the number of ASVs (a), Shannon (b) and inverse Simpson indices (c). Lowercase letters indicate significant differences between treatments (using ANOVA followed by Tukey posthoc tests, p ≤ 0.05).

PCoA ordination analyses performed on the entire microbial community data (based on the Bray-Curtis distance) showed that samples separated into four groups according to salinity and experimental treatments (Figure 5; ANOSIM p < 0.001), where some dispersal appeared between sampling days (for the field campaigns). This implies that the microbial community in Achziv porewater is relatively stable annually and diversity is primarily influenced by water salinity. The low-salinity porewater (n = 9) was the most distinctive compared to the other groups of the high-salinity porewater (n = 13), Achziv surface water (n = 11), and the experimental treatments: non-filtered experimental treatment (n = 11) and seawater control group (n = 4). The exceptions were the 5% 0.1 µm filtered treatment (n = 3; p > 0.05) and fresh groundwater samples (n = 3; p > 0.05). Overall, the PCoA plot was divided into four main quarters along both axes, each representing different water characteristics (Figure 5). While, the left half of the PCoA2 axis contained samples originating from fresh groundwater and low-salinity porewater, the right half was dominated by high-salinity porewater and surface seawater, including samples from Achziv and the experimental treatments. During the field campaign in February 2021, two surface seawater samples were collected at different distances from the shore (100m and 600m). It is apparent from Figure 5 that the high-salinity porewater community composition in the aquifer was more similar to that of the near-shore (100m) Achziv surface seawater than to that of the open seawater (600m from the shore). These communities possibly affect each other, creating a new environment across the seawater-aquifer transect. For the same field sampling campaign, we measured surface seawater and porewater microbial activity (primary and heterotrophic production) and prokaryotic abundance. We observed a substantial decrease in all parameters between the conservative mixing line and seawater (Supplementary data, S5). These findings illustrate the change in the environment between open seawater and groundwater, which both prevents day light and introduces new communities, which may limit productivity.

**Figure 5:**
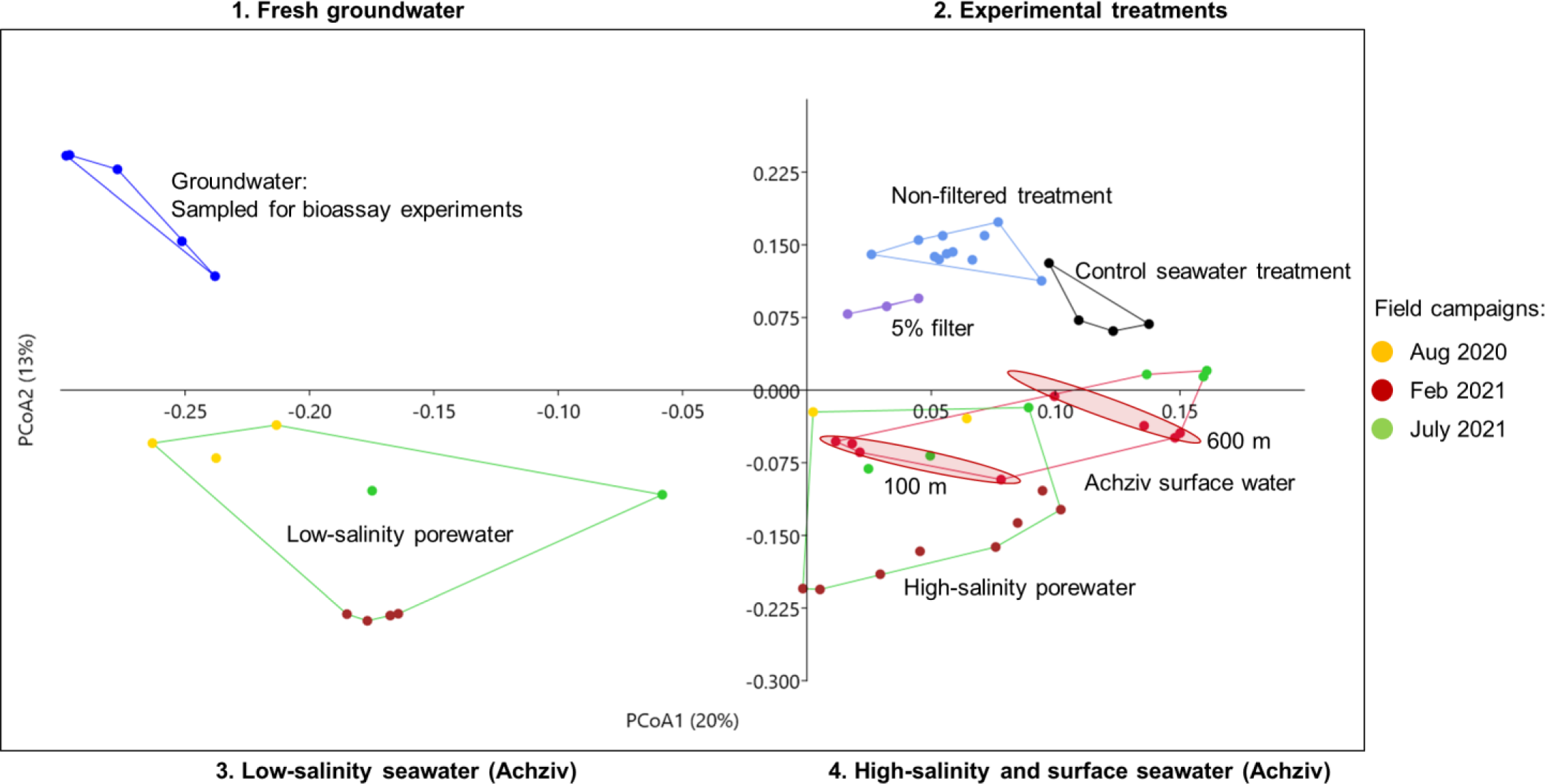
Principal coordinate analysis (PCoA) of prokaryote composition (Bray-Curtis distances) based on 16S-amplicon sequencing of samples collected during three sampling campaigns (different colors as indicated) at Achziv site, grouped according to porewater salinity (Low/High) and Surface seawater. The Achziv surface seawater samples collected in February 2021 were divided according to distance from shore (100 and 600m). Additionally, experimental treatment samples (control seawater, non-filtered, 5% filtered and fresh groundwater) are plotted.

Achziv bacterial community composition (surface seawater) was co-dominated by three marine family-level taxonomic groups (relative abundance > 40%), including the autotrophic *Synechococcus* and heterotrophs (Figure 6, Supplementary Table S1). The three heterotrophic bacterial families were *Rhodobacteraceae*, *Actinomarinaceae* and SAR11 Clade. While *Actinomarinaceae* was the most abundant bacterial plankton in all surface seawater samples, including the experimental control seawater (ambient seawater, not influenced by SGD), Achziv surface seawater and high-salinity porewater, the SAR11 clade was primarily abundant in the control seawater and non-filtered experimental treatments, Figure 6a&b, respectively. On the other hand, the lowest relative abundance of *Rhodobacteraceae* was detected in experimental control seawater, comparable with fresh groundwater samples. Thus, from ambient coastal seawater to SGD-coastal samples (Achziv), the community composition shifted from a SAR11-dominated to a *Rhodobacteraceae*-dominated community. Notably, the relative abundance of *Rhodobacteraceae* and *Alteromonadaceae* were the highest in the filtered treatment (Figure 6c & d, respectively). Previous studies have linked the presence of these marine microbial taxa in coastal aquifers as potential indicators of seawater intrusion, indicating an active land-ocean interface (Chen et al., 2019; Unno et al., 2015).

**Figure 6:**
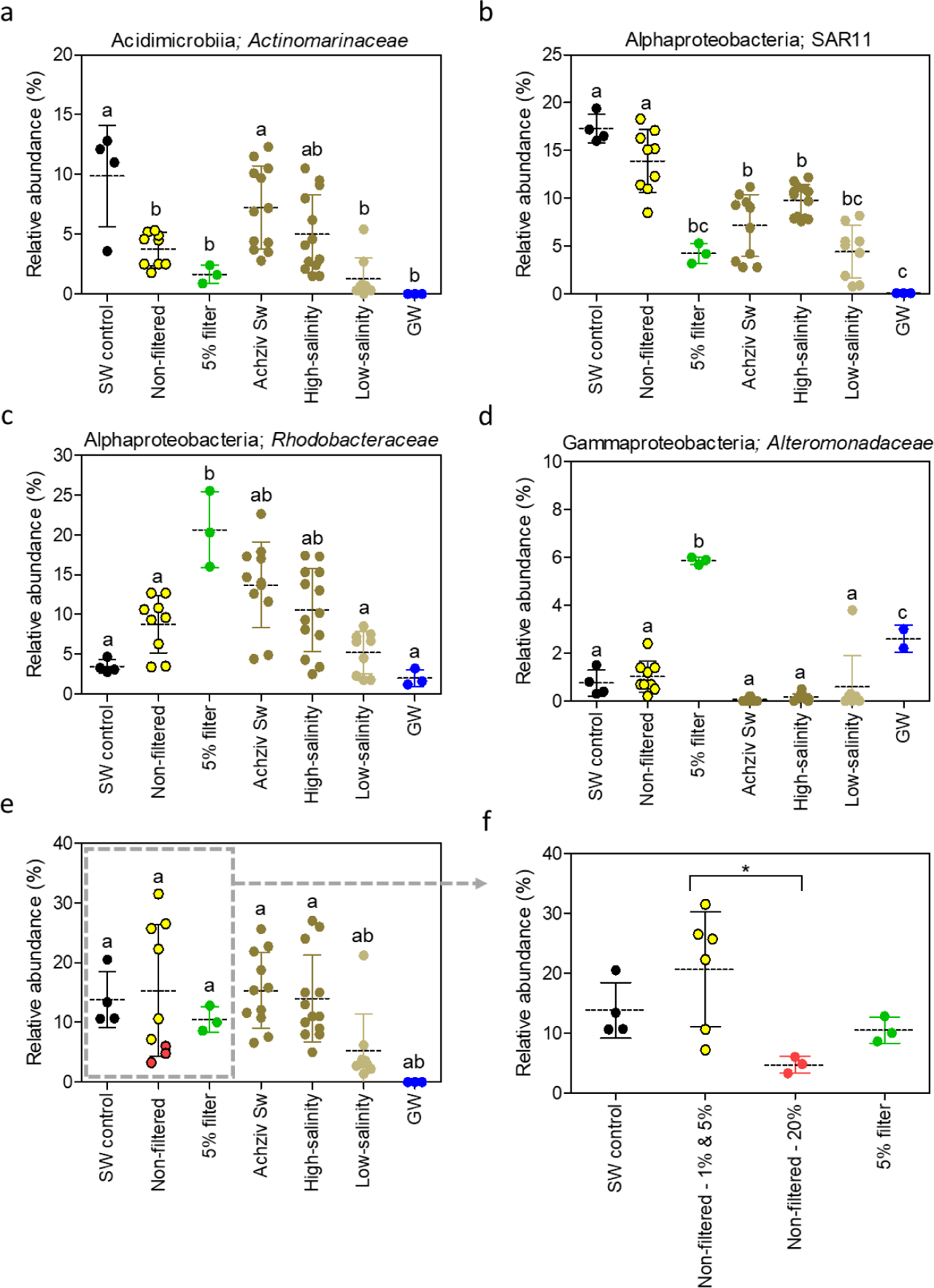
Scatter dot plots of the four dominant bacterial groups (relative abundances): (a) *Actinomarinaceae* (b) SAR11 (c) *Rhodobacteraceae* (d) *Alteromonadaceae* (e) the autotrophic bacterium *Synechococcus* in Achziv field samples (samples grouped according to water-salinity) and incubation experiments (samples grouped according to treatment). (f) *Synechococcus* relative abundances in incubation samples, differentiating between the different treatments. Lowercase letters indicate significant differences between treatments (using ANOVA followed by Tukey posthoc tests, p ≤ 0.05). * indicates significant difference between two treatments (p < 0.05), where the rest are non-significant. Data derived from 16S rDNA analyses.

Based on the SAR11 to *Rhodobacteraceae* shift and the substantially higher relative abundance of *Rhodobacteraceae* in the filtered treatment, we suggest that applying the filtration resulted in a unique scenario, where the typical-oligotrophic coastal microbial community, growing under constant nutrient-limiting conditions, was exposed to a nutrient-rich solution almost without potential competitors.

Our results suggest that mixing non-filtered groundwater with ambient coastal seawater induced competitive interactions between the two different communities, eventually affecting primary and heterotrophic production rates. This competitive effect was essentially eliminated through filtration because most microbes were removed from the water, relieving the nutrient limitations of the oligotrophic coastal seawater. Although the transport of microbial diversity into the coastal ocean through SGD has been addressed mostly with regard to pathogenic or fecal indicator bacteria (Vollberg et al., 2019; Yau et al., 2014), our results emphasize that marine heterotrophic bacterioplankton respond taxonomically and functionally to SGD-derived microbes. Typically, low-salinity porewater and fresh groundwater samples were similar to each other in terms of low relative abundance of the marine bacteria found in the other samples (Figure 6a, b, c) and β-diversity distance (ANOSIM, p > 0.05). Moreover, the higher alpha diversity values in the low-salinity porewater samples (Figure 4) are explained by a much more heterogeneous microbial community, which consisted of a diverse mixture of freshwater and marine microbes. These include the prevalent groundwater clades Patescibacteria and Woesearchaeales (Herrmann et al., 2019; Tian et al., 2020). Patescibacteria was found in low abundances in Achziv surface seawater, and absent in the control ambient samples (Supplementary data, Table S2). This hints at the potential role of connectivity between terrestrial and marine hydrospheres driven by SGD in the Achziv site.

*Synechococcus* was the dominant cyanobacterium detected in all surface seawater and high-salinity porewater samples (Figure 6e). The lowest abundance was associated with the low-salinity porewater and groundwater samples, probably due to the different environmental conditions, primarily the lack of light in the aquifer. Moreover, the highest groundwater addition treatment (20%) had the lowest *Synechococcus* relative abundance when compared to the other lower ratios of non-filtered treatments (Figure 6f), suggesting larger volumes of groundwater inhibit *Synechococcus* growth. However, this was less significant based on cell number counts (Supplementary Table S1).

The third experiment (hereafter Exp. 3) was designed to extend on the previous two experiments and further examine the response of autotrophic communities to different microbial cell size fractions by mixing sequentially filtered groundwater (non-filtered, 0.22 & 0.1 µm pore size) with the control ambient seawater, Figure 7. Also here, heterotrophic production and prokaryote abundance increased when 0.1 µm filtered groundwater was mixed with the control seawater. However, the 0.22 µm filtration treatment did not exert such a response on prokaryote abundance and heterotrophic production, rather induced primary productivity rates, specifically associated with *Synechococcus* abundance, after a 40-hour incubation (Figure 7a&b). Cell abundances dropped for all treatments after a 65-Hr incubation period, but even the delayed production rates measured only for the last time period (65Hr) were still showing a significant response, which supported the abundance trend observed after a 40-Hr incubation period. The DDW treatment was included to demonstrate that groundwater mixing effects occur as a result of terrestrial-coastal microbial interactions, and not due to simple volumetric dilution of the seawater with fresh water.

**Figure 7:**
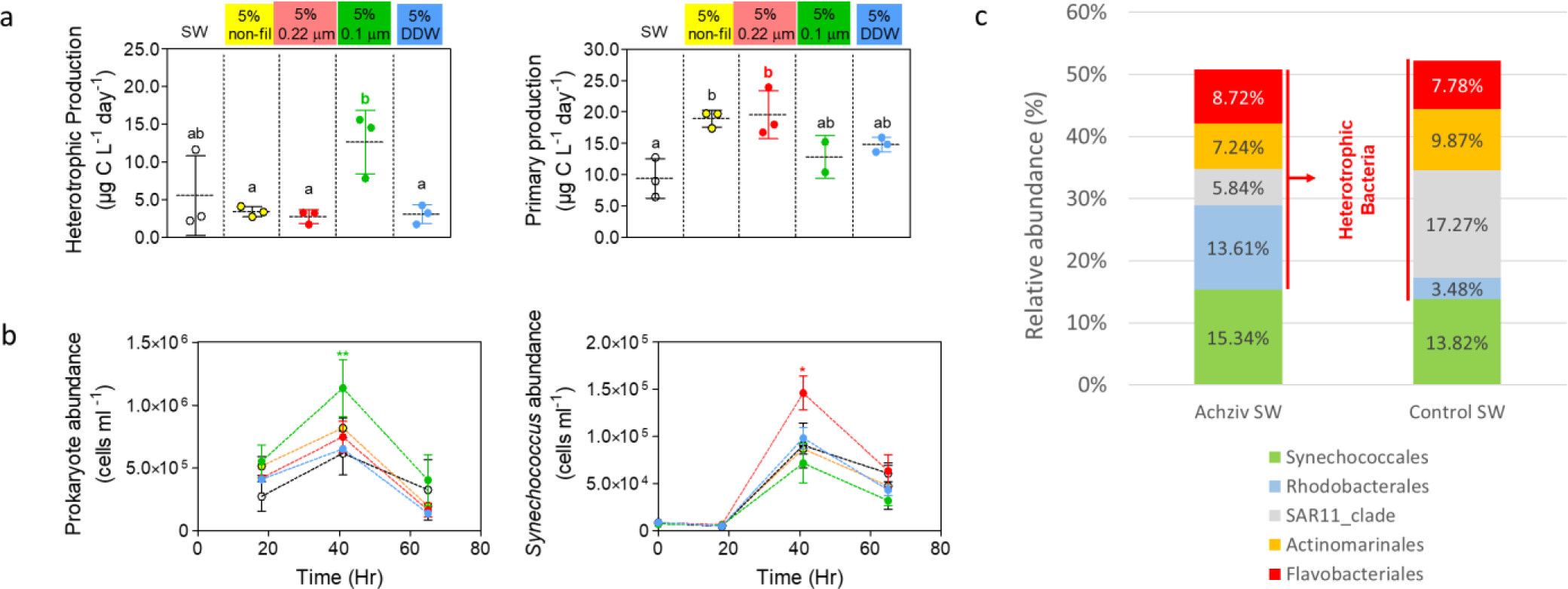
a) Scatter dot plots of heterotrophic and primary production rates following dilution with fresh groundwater (5%, 5% filtered 0.22 μm and 5% filtered 0.1 μm v:v), un-amended seawater (SW) or diluted with 5% distilled water (5% DDW) during Experiment 3. Results are shown for the final time point of the experiment (only initial and final measurements were conducted for this experiment). Lowercase letters indicate significant differences between treatments (using ANOVA followed by Tukey posthoc tests, p ≤ 0.05). b) Temporal variability of prokaryote and *Synechococcus* abundance detected (data derived through flow cytometry); */** indicate significant difference between two treatments (p < 0.05 or p < 0.01, respectively), where the rest are non-significant. c) Bar plot showing relative sequence abundance of dominant microbial taxa at order level for both coastal sites

### 4.5 SGD affects the interplay between *Synechococcus* and heterotrophic bacteria

In our two investigated sites (SGD and control), *Synechococcus* was the dominant cyanobacterium based on 16S data. Although *Synechococcus* relative abundance was similar for both sites, we observed marked differences in the composition of heterotrophic bacteria, specifically SAR11 and *Rhodobacteraceae* groups (Figure 7c). The strict oligotrophic SAR11 clade are highly abundant in the open sea, and were found to associate with Cyanobacteria primarily under poor dissolved organic carbon and nutrient conditions (Kearney et al., 2021). Alternatively, *Rhodobacteraceae* is much more metabolically flexible (Xia et al., 2021), allowing this group to outcompete and dominate other heterotrophs upon groundwater discharge.

Therefore, heterotrophic bacterioplankton largely determine carbon biogeochemical cycling in the ocean through respiration and biomass production, and their composition appears to be affected by SGD (Kaile’a and Wiegner, 2016). These results suggest that *Synechococcus* abundance can increase similarly in the two sites (SGD-enriched and oligotrophic), but primary production activity is limited, maintaining a balanced ecological ecosystem (perhaps as an intrinsic control of harmful-algal blooming). Currently, the mechanistic role SGD plays in maintaining ecosystem stability remains unknown, and how the groundwater microbiota drives this effect, particularly concerning different microbial cell size fractions.

### 4.6 potential effect on coastal heterotrophic autotrophic balance and carbon cycling in the ocean

Although freshwater SGD was found to be minor compared to freshwater river fluxes to the ocean (Luijendijk et al., 2020; Zhou et al., 2019), saline and brackish water fluxes are much more significant (e.g. Beck et al., 2013; Kwon et al., 2014; Santos et al., 2021). Our results show how SGD shape the heterotrophic community diversity and regulate activity through discharged microbes. These observations have significant implications for the coastal and ocean carbon budget. Carbon fixation by autotrophs and heterotroph respiration shape the carbon budget in the ocean, affecting directly atmospheric CO_2_ through the ability to bury organic carbon (Kirchman, 2012). Estimating chemolithoautotroph fixation rates in STE ecosystems are also important given the notable inorganic carbon fixation rates recently reported in a carbonate aquifer (Overholt et al., 2022). Climate change, causing global warming and ocean acidification have shown to enhance heterotrophic activity (Duarte et al., 2013), which may affect the balance between autotrophic carbon fixation and heterotrophic recycling, thus the biogenic carbon fluxes in the ocean. Our study suggests that this relationship may be more complex through changes in SGD fluxes due to sea level changes. An increase in sea level will reduce SGD fluxes (Kiro et al., 2008), and thus affecting the heterotrophic community in seawater, as concluded in this study. This may have direct implications on the delicate balance between heterotrophs and autotrophs, and the biogenic carbon fluxes.

## 5 Conclusions

SGD is an important source of nutrients in coastal environments, especially in oligotrophic environments. In this study we shed light on the complex contribution of SGD on the ecology of coastal microbial communities through the transport of groundwater-borne prokaryotes. Diverse microbial communities, naturally inhabiting the subterranean estuary, shape coastal microbial abundance, taxonomic diversity and activity. This ultimately affects photoautotrophic-heterotrophic specific interactions, marine food webs and biogeochemical cycles of the entire coastal ecological system. In our current changing climate world, these interactions are crucial for understanding the possible effects and feedback mechanisms on carbon cycling and climate. Further research is needed to integrate the global chemical-biological effect of SGD on marine environments, focusing, for example, on STE sediment microbial dynamics along redox gradients at the terrestrial–marine interface.

## Acknowledgments

We thank to the Sustainability and Energy Research Initiative (SAERI) grant, the Helen Kimmel Center for Planetary Science grant and the De Botton center for Marine Sciences grant for their funding supporting this study. We also thank Dr. Daniella Gat for her help and advice in bioinformatics analysis. This study was supported by the, Zuckerman Faculty Scholars program, program, a research grant from the Center for Scientific Excellence, a research grant from the Raymond Lapon Fund, and the Estate of David Levinson, a research grant from Paul and Tina Gardner, and a research grant from the Center for New Scientists at the Weizmann Institute of Science. We thank H. Michael and I. Halevy for fruitful discussions and suggestions.

