## Supporting Information for "Allochthonous groundwater microorganisms affect coastal seawater microbial abundance, activity and diversity"

### Contents of this file

Figures S1 to S5
Table S1 and Table S2

### Introduction

This Supporting Information contains five additional figures and two additional tables on microbial abundance and activity, nutrient concentration changes during experiment 2, and mixing lines of porewater and surface seawater measurements vs. water density collected from Achziv beach. Table S1 summarizes activity, abundance and community composition of all samples collected during the study (three incubation experiments and three field campaigns). Table S2 compares community composition of the fresh groundwater and low-salinity porewater samples.

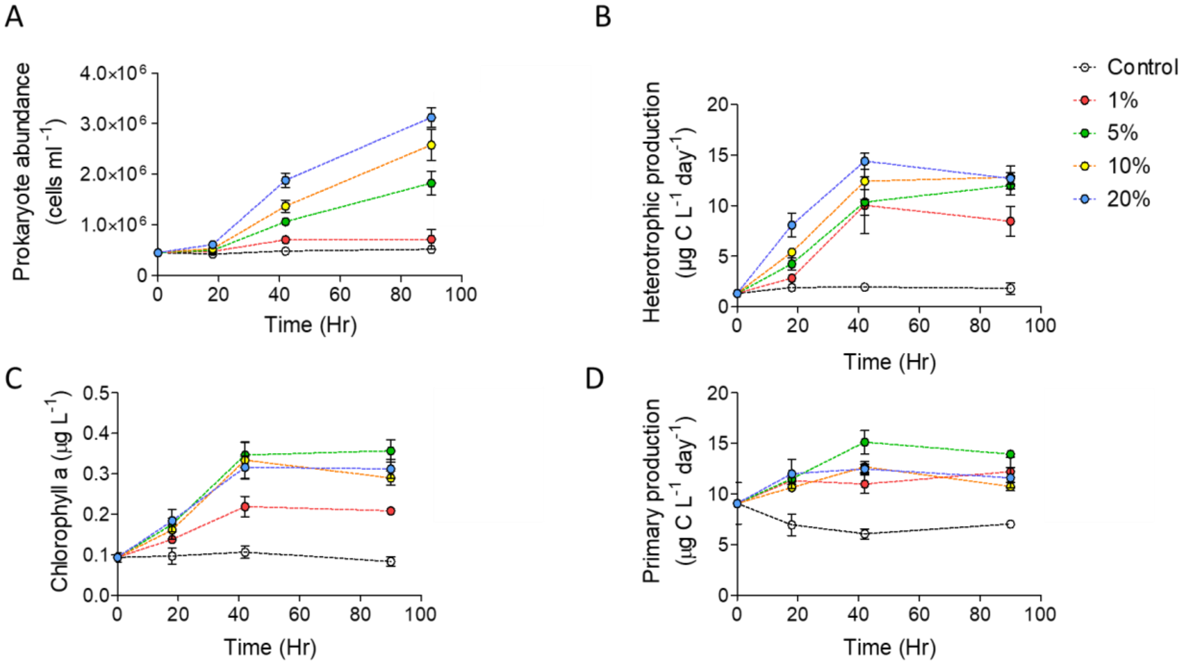


**Figure S1.** Temporal variability of surface seawater (control) prokaryote abundance (A), heterotrophic production rate (B) primary production rate (C) and Chlorophyll a concentrations (D) following dilution with discharged brackish groundwater (1-20% v:v) or un-amended seawater (control) during Experiment 1. The dilution factor was calculated for each treatment to account for the volume of groundwater added to ambient seawater.

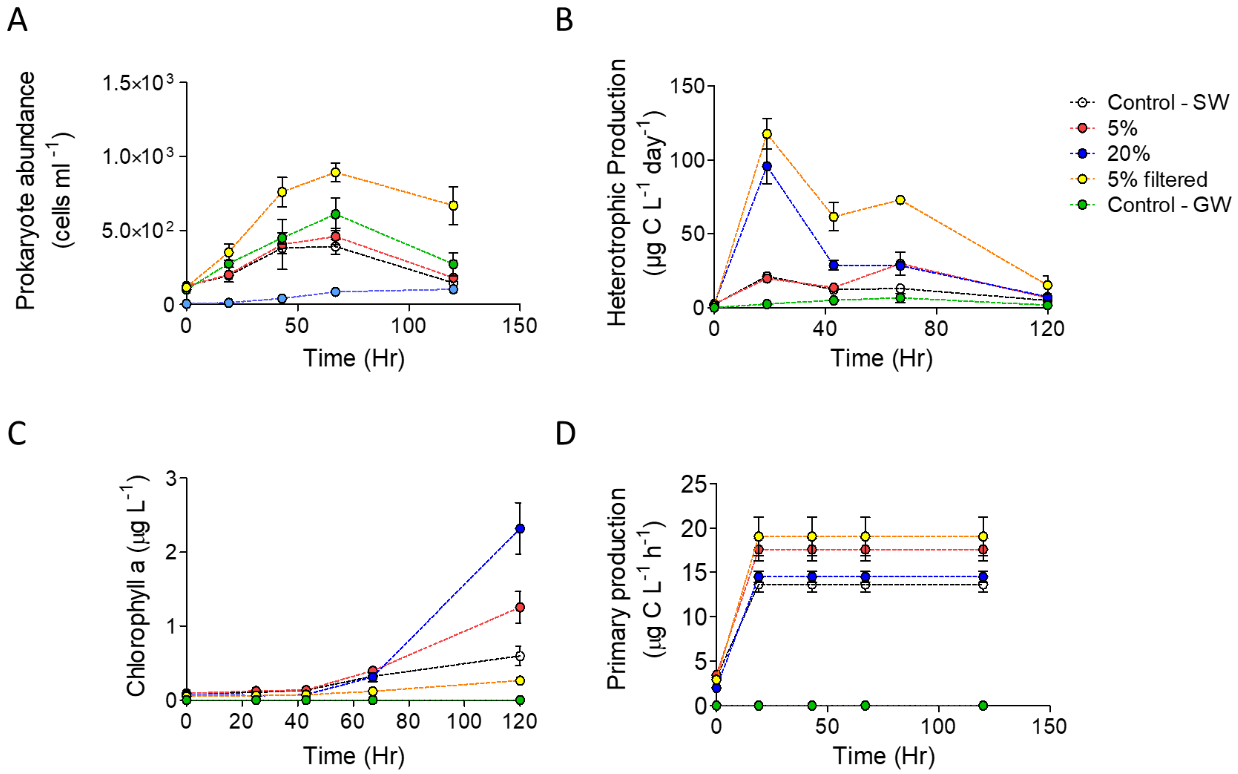


**Figure S2.** Temporal variability of surface seawater (control) prokaryote abundance (A), heterotrophic production rate (B) primary production rate (C) and Chlorophyll a concentrations (D) following dilution with fresh groundwater (5%, 20% and 5% filtered 0.1 µm v:v), un-amended seawater (control SW) or fresh groundwater (Control GW) during Experiment 2. The dilution factor was calculated for each treatment to account for the volume of groundwater added to ambient seawater.

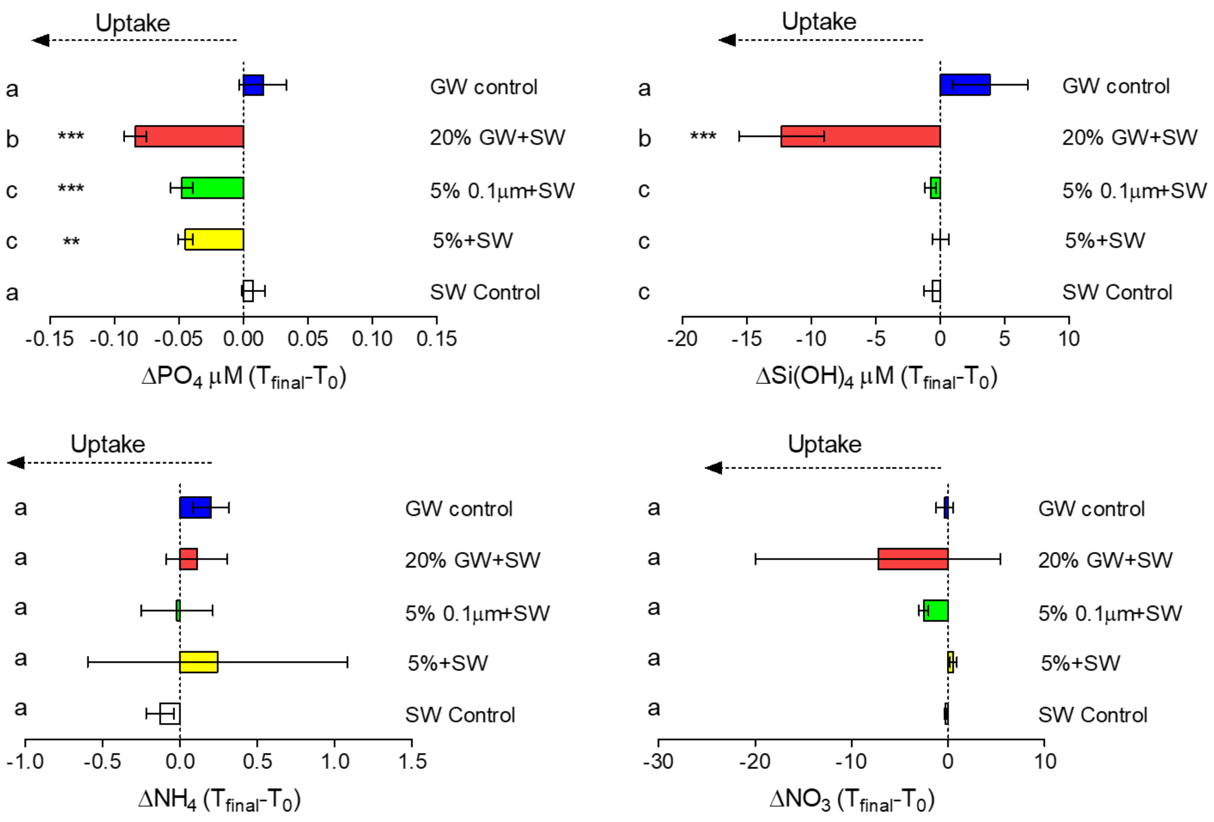


**Figure S3.** Change in phosphate, silicate, ammonium and nitrate concentration in the water of each treatment over the 5-day interval of experiment 2. Negative values indicate microbial uptake. Units are in µM. Lowercase letters indicate significant differences between treatments (using ANOVA followed by Tukey posthoc tests, $p \leq 0.05$).

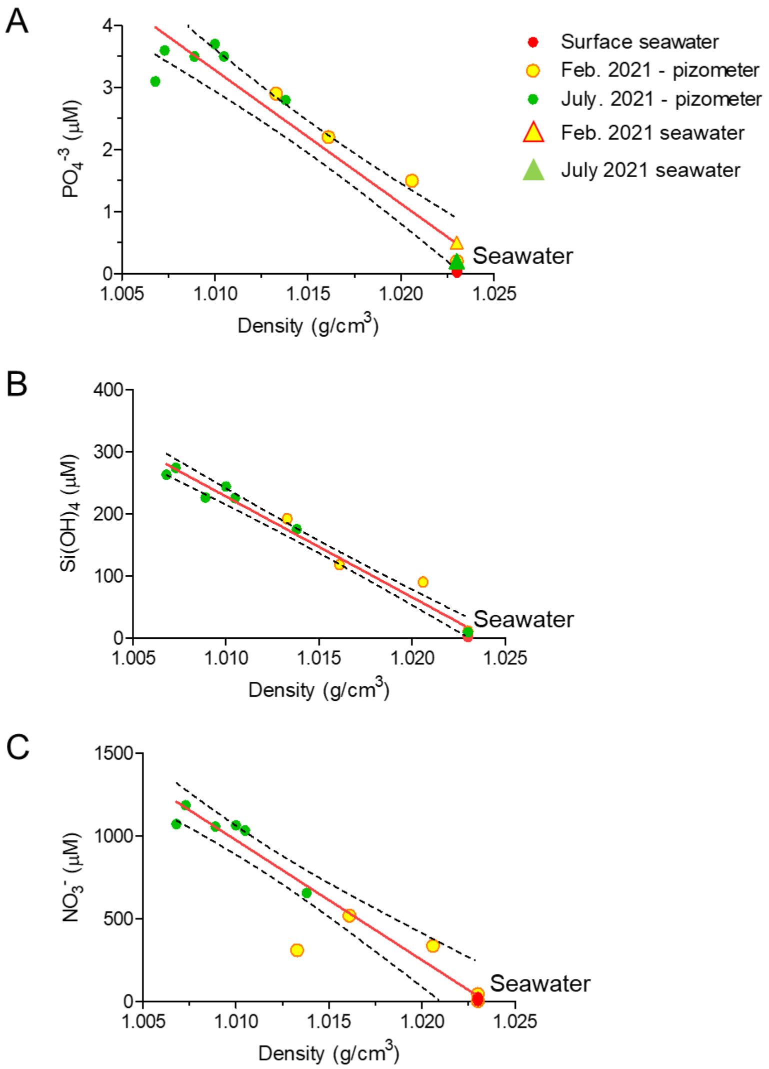


**Figure S4.** (A) $PO_4^{-3}$ (B) $Si(OH)_4$ (C) $NO_3^-$ vs. water density in porewater using piezometers (Figure 1, main text) and open seawater from Achziv beach.

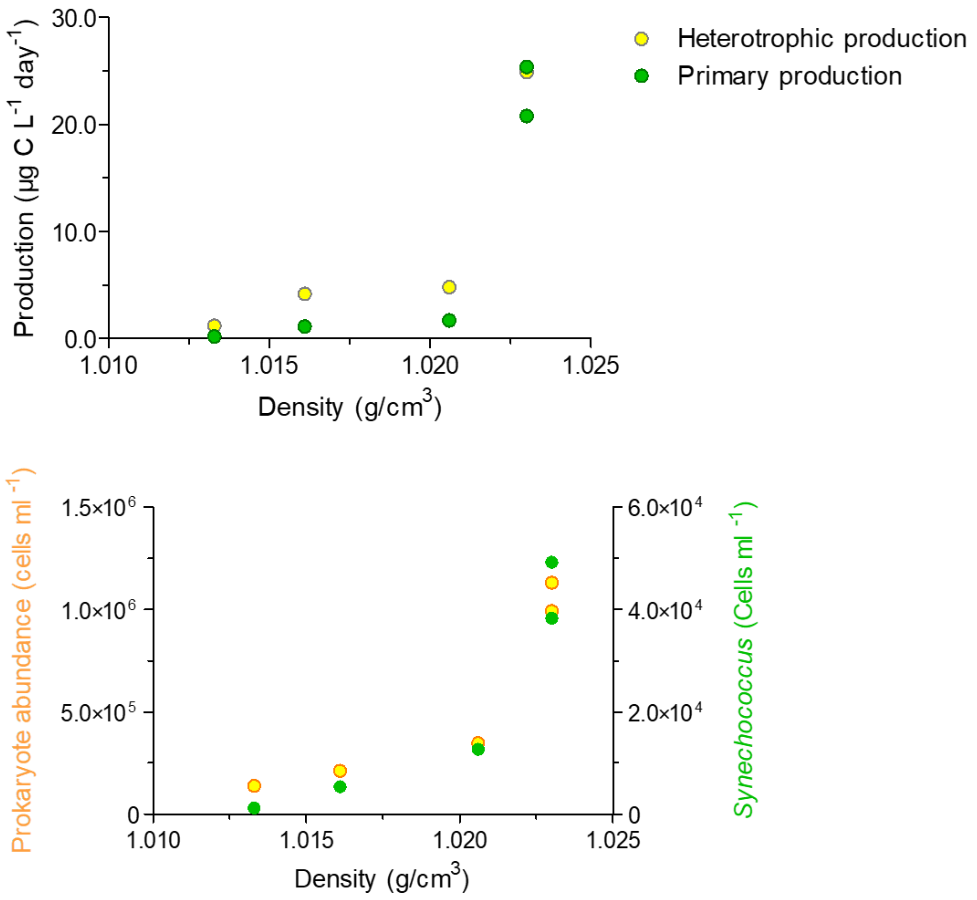


**Figure S5:** Microbial production (top) and abundance (bottom) vs. water density in porewater using piezometers (Figure 1, main text) and open seawater from Achziv beach.

**Table S1**: Summary of community composition, activity and microbial abundance averaged by sample groups

| Sample source | Group | Relative abundance (%) of four marine dominant families | | | | | Abundance and activity [a, b] | | | |
|---|---|---|---|---|---|---|---|---|---|---|
| | | *Synechococcus* Cyanobacteriia | *Rhodobacteraceae* Alphaproteobacteria | *Actinomarinaceae* Acidimicrobiia | SAR11 Clade Alphaproteobacteria | *Alteromonadaceae* Gammaproteobacteria | Prokaryote abundance (Cells/ml) | Heterotrophic production (µg C $L^{-1}$ $day^{-1}$) | *Synechococcus* abundance (Cells/ml) | Primary production (µg C $L^{-1}$ $day^{-1}$) |
| Environmental Achziv study site | Low-salinity porewater | 5.2± 6.1 | 5.2± 2.6 | 1.3± 1.7 | 4.4±2.7 | 0.6±1.3 | 1.3x $10^5$ | 1.2±0.2 | 1.2x$10^3$ | 0.2± 0.09 |
| | High-salinity porewater | 14.0± 7.2 | 10.6± 5.2 | 5.0± 3.3 | 9.8±1.7 | 0.2±0.1 | 2.8x $10^5$ | 4.5±2.2 | 9.0x$10^3$ | 1.4±0.3 |
| | Achziv surface seawater | 15.3± 6.3 | 13.7± 5.4 | 7.2± 3.5 | 7.2±3.2 | 0.05± 0.08 | 1.0x $10^6$ | 29.3± 6.3 | 3.7x$10^4$ | 27.0± 7.1 |
| Incubation experiments | Control seawater | 13.8± 4.6 | 3.5± 0.8 | 9.9± 4.2 | 17.3± 1.5 | 0.75± 0.5 | 4.9x $10^5$ | 6.8±5.6 | 3.7 x$10^4$ | 9.7±3.7 |
| | Non-filtered: | **1-5%:** 20.6± 9.6 **20%:** 4.7±1. 4 | 8.8± 3.6 | 3.8± 1.4 | 13.9± 3.2 | 1.02± 0.6 | 9.9x $10^5$ | 17.6± 9.0 | **1-5%:** 2.5 x$10^4$ **20%:** 2.0 x$10^4$ | 14.6± 2.8 |
| | 5%-filtered | 10.5± 2.1 | 20.6± 4.7 | 1.6± 0.7 | 4.2±1.1 | 5.9±0.1 | 1.02 x $10^6$ | 42.8± 33.1 | 7.3 x$10^4$ | 16.6± 4.1 |
| | Fresh groundwater | 0.0± 0.0 | 2.0± 1.1 | 0.0± 0.0 | 0.1±0.0 | 2.6±0.6 | 4.5 x $10^5$ | 8.5±2.9 | 1.8 x$10^2$ | 0.2±0.3 |

[a] Results for abundance and activity of field samples were measured for one campaign (February 2021).

[b] Bottle incubation results include all values measured for experiments 1-3 after 40-72 Hr incubation time.

**Table S2**: Relative abundance of low-salinity and fresh groundwater community composition.

**Control GW**

| Class | Order | Relative abundance |
|---|---|---|
| Firmicutes | Bacilli | 14.5% |
| Alphaproteobacteria | Rhodospirillales | 15.2% |
| Alphaproteobacteria | Caulobacterales | 8.3% |
| Alphaproteobacteria | Sphingomonadales | 26.9% |
| Alphaproteobacteria | Rhodobacterales | 5.7% |
| Alphaproteobacteria | Rhizobiales | 3.5% |
| Gammaproteobacteria | Alteromonadales | 4.9% |
| Gammaproteobacteria | Oceanospirillales | 10.9% |
| Gammaproteobacteria | Burkholderiales | 2.7% |
| Gammaproteobacteria | Pseudomonadales | 1.5% |
| Gammaproteobacteria | Legionellales | 1.8% |
| Desulfuromonadia | Bradymonadales | 2.4% |
| Omnitrophia | Omnitrophales | 1.5% |

**Low-salinity**

| Class | Order | Relative abundance |
|---|---|---|
| Cyanobacteriia | Synechococcales | 5.2% |
| Bdellovibrionia | Bdellovibrionales | 8.0% |
| Oligoflexia | | 2.1% |
| Alphaproteobacteria | Rhodobacterales | 5.9% |
| Alphaproteobacteria | SAR11_clade | 5.3% |
| Alphaproteobacteria | Rhizobiales | 2.7% |
| Alphaproteobacteria | Rhodospirillales | 0.8% |
| Alphaproteobacteria | Sphingomonadales | 1.4% |
| Gammaproteobacteria | Oceanospirillales | 1.3% |
| Gammaproteobacteria | Alteromonadales | 3.2% |
| Gammaproteobacteria | Burkholderiales | 0.6% |
| Gammaproteobacteria | Cellvibrionales | 0.7% |
| Gammaproteobacteria | Oceanospirillales | 1.3% |
| Gammaproteobacteria | Vibrionales | 0.7% |
| Desulfobulbia | Desulfobulbales | 1.2% |
| Desulfuromonadia | Bradymonadales | 1.0% |
| Patescibacteria | Gracilibacteria | 2.7% |
| Patescibacteria | ABY1 | 3.8% |
| Patescibacteria | Parcubacteria | 4.1% |
| Patescibacteria | unclassified | 10.7% |
| Lentisphaeria | Lentisphaerales | 0.8% |
| Nanoarchaeia | Woesearchaeales | 5.5% |
